# Hierarchical sparse coding of objects in deep convolutional neural networks

**DOI:** 10.1101/2020.06.29.176032

**Authors:** Xingyu Liu, Zonglei Zhen, Jia Liu

## Abstract

Recently, deep convolutional neural networks (DCNNs) have attained human-level performances on challenging object recognition tasks owing to their complex internal representation. However, it remains unclear how objects are represented in DCNNs with an overwhelming number of features and non-linear operations. In parallel, the same question has been extensively studied in primates’ brain, and three types of coding schemes have been found: one object is coded by entire neuronal population (distributed coding), or by one single neuron (local coding), or by a subset of neuronal population (sparse coding). Here we asked whether DCNNs adopted any of these coding schemes to represent objects. Specifically, we used the population sparseness index, which is widely-used in neurophysiological studies on primates’ brain, to characterize the degree of sparseness at each layer in two representative DCNNs pretrained for object categorization, AlexNet and VGG11. We found that the sparse coding scheme was adopted at all layers of the DCNNs, and the degree of sparseness increased along the hierarchy. That is, the coding scheme shifted from distributed-like coding at lower layers to local-like coding at higher layers. Further, the degree of sparseness was positively correlated with DCNNs’ performance in object categorization, suggesting that the coding scheme was related to behavioral performance. Finally, with the lesion approach, we demonstrated that both external learning experiences and built-in gating operations were necessary to construct such a hierarchical coding scheme. In sum, our study provides direct evidence that DCNNs adopted a hierarchically-evolved sparse coding scheme as the biological brain does, suggesting an implementation-independent principle of representing a myriad of objects efficiently.

## 1 Introduction

One spectacular achievement of human vision is that we can accurately recognize objects at a fraction of a second in the complex visual world. In recent years, deep convolutional neural networks (DCNNs) have achieved human-level performances in object recognition tasks (He et al., 2015; Simonyan and Zisserman, 2015; Szegedy et al., 2015). The success is primarily credited to the architecture that generic DCNNs compose of a stack of convolutional layers and fully-connected layers, each of which has multiple units with different filters (i.e., ‘neurons’ in DCNNs), similar to the hierarchical organization of primates’ ventral visual stream. With such hierarchical architecture and supervised learning on a large number of object exemplars, DCNNs are thought to construct complex internal representations for external objects. However, little is known about how exactly objects are represented in DCNNs.

This question has already puzzled neuroscientists for a long time. To understand how primates’ visual system encodes external world, three types of coding schemes are proposed to describe how neurons are integrated together to represent an object. At one extreme is distributed coding, by which the whole neuronal population is involved, whereas at the other extreme is local coding, by which one neuron is designated to represent one object. The distributed coding scheme is superior in large coding capacity, easy generalization and high robustness, while the local coding scheme is good at information compression, energy conservation and better interpretability. In between lies the sparse coding that different subsets of neurons in the population participate in coding different objects. As a trade-off, sparse coding possesses advantages of both local coding and distrusted coding (Barlow, 1972; Thorpe, 1989; Berkes et al., 2009; Rolls, 2017; Beyeler et al., 2019; Thomas and French, 2017). Neurophysiological studies have revealed that the sparse coding scheme is adopted in some areas in primate visual cortex for object recognition (Olshausen and Field, 1996; Lehky et al., 2011; Barth and Poulet, 2012; Rolls, 2017).

Following the studies on biological intelligent system, several pioneer studies started to characterize DCNNs’ representation with coding scheme (Szegedy et al., 2013; Agrawal et al., 2014; Li et al., 2016; Wang et al., 2016; Morcos et al., 2018; Casper et al., 2019; Parde et al., 2020). Studies using the ablation approach shows that the processing of objects usually requires the participation of multiple units, but only 10% - 15% of units in a layer are actually needed to achieve 90% of the full performance (Agrawal et al., 2014). Even when half of the units in all layers are ablated, the performance does not decrease significantly with the accuracy above 90% of the full performance (Morcos et al., 2018). Further studies quantify the number of non-zero units in response to objects and report a trend of decrease in number of non-zero units along the hierarchy of DCNNs(Agrawal et al., 2014). These preliminary results suggest that DCNNs may adopt the sparse coding scheme, which likely evolves along hierarchy.

Here, we adopted a prevalent metric in neurophysiological studies on primates’ brain, population sparseness index (PSI, Rolls and Tovee, 1995; Vinje and Gallant, 2000), to quantify the population sparseness along the hierarchy of two representative DCNNs, AlexNet (Krizhevsky, 2014) and VGG11 (Simonyan and Zisserman, 2015). Specifically, we first systematically evaluated the layer-wise sparseness in representing objects. Then, we characterized the functionality of sparseness by examining the relationship between sparseness and behavioral performance in each layer. Finally, we explored factors that may influence the coding scheme.

## 2 Materials and methods

### 2.1 Visual images datasets

#### ImageNet dataset

The dataset from ImageNet Large Scale Visual Recognition Challenge 2012 (ILSVRC2012, http://www.image-net.org/) (Russakovsky et al., 2015) contains 1,000 categories that are organized according to the hierarchy of WordNet (Miller, 1995). The 1,000 object categories consist of both internal nodes and leaf nodes of WordNet, but do not overlap with each other. The dataset contains 1.2 million images for model training, 50,000 images for model validation and 100,000 images for model test. In the present study, only the validation dataset (i.e., 1,000 categories × 50 images) was used to evaluate the coding scheme of DCNNs.

#### Caltech256 dataset

The Caltech256 dataset consists of 30,607 images from 256 object categories with a minimum number of 80 images per category. In the present study, 80 images per category were randomly chosen from the original dataset.

### 2.2 DCNNs and activation extraction

The well-known AlexNet and VGG11 that are pretrained for object classification were selected to explore the coding scheme of DCNNs. Besides the two pretrained models, corresponding weight-permuted models and ReLU-deactivated models were also examined to investigate the factors that may influence the coding scheme observed in the pretrained models.

#### Pretrained models

AlexNet and VGG11 are pretrained on ILSVRC2012 dataset and were downloaded from PyTorch model Zoo (https://pytorch.org/). Both DCNNs are purely feedforward: the input to each layer consists solely of the output from the previous layer. The AlexNet consists of 5 convolutional layers (Conv1 through Conv5) that contain a set of feature maps with linear spatial filters, and 3 fully-connected layers (FC1 through FC3). In between, a max (x, 0) rectifying non-linear unit (ReLU) is applied to all units after each convolutional and FC layer. In some convolutional layers, ReLU is followed by another max-pooling sublayer. VGG11 is similar to AlexNet in architecture except for two primary differences. First, VGG11 uses smaller receptive fields (3×3 with a stride of 1) than AlexNet (11×11 with a stride of 4). Second, VGG11 has more layers (8 convolutional layers) than AlexNet. When we refer to Conv#, we mean the outputs from the ReLU sublayer in the #th convolutional layer. Similarly, FC# means the outputs from the #th FC layer after ReLU. The DNNBrain toolbox (https://github.com/BNUCNL/dnnbrain/) was used to extract the DCNN activation. The activation from all elements within a unit (or channel) was averaged to produce a unit-wise (or channel-wise) activation for each exemplar, and the activation of the unit to an object category was then derived by averaging the unit-wise responses from all exemplars of the category.

#### Weight-permuted models

The weight permuted model was derived by permuting weights of the pretrained models within each layer. That is, the structures of the original networks and the weight distribution of each layer were preserved while the exact feature filters obtained from the learning of the supervised task were disrupted.

#### ReLU-deactivated models

The ReLU-deactivated model was the same with the pretrained models with only ReLU being silenced in all layers by replacing it with an identity mapping. The ReLU-deactivated model disabled the non-linear operation after the feature extraction but still retained the same network architectures and the learned feature filters.

### 2.3 Population sparseness index

The PSI was calculated for each layer of DCNNs to quantify the peakedness of the distribution of population responses elicited by an object category, which is equivalent to the fraction of the units in the population that participated in coding objects in the case of binary responses (Vinje and Gallant, 2000).

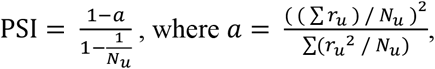

where r_*u*_ is the unit-wise activation of a unit u from a target layer in response to an object category, and N_*u*_ is the number of units in that layer. The unit-wise activation was z-scored across all categories for each unit, and then normalized across all units into a range from 0 to 1 to rescale the negative values to non-negative as required by the definition of PSI. Values of PSI near 0 indicate low sparseness that all units respond equally to the object category, and values near 1 indicate high sparseness that only a few units respond to the category.

### 2.4 Relationship between population sparseness and classification performance

The relationship between sparseness and classification performance was first explored using correlation analyses. The Caltech256 classification task was used to estimate the classification performance of AlexNet and VGG11 on each category. Specifically, a logistic regression model was constructed using activation patterns from FC2 as features to perform a 256-class object classification. A 2-fold cross-validation procedure was used to evaluate the classification performance. Then, Pearson correlation coefficients between the PSI and the classification performance were calculated across all categories for each layer respectively. Finally, to reveal how the sparse coding from different layers contribute to the classification performance, a stepwise multiple regression was conducted with the classification performance of each category as dependent variables and the PSI of the corresponding category from all layers as independent variables. The regressions were conducted for Conv layers and FC layers separately.

## 3 Results

The coding scheme for object categorization in DCNN was characterized layer by layer in the pretrained AlexNet and VGG11 using PSI. The PSI was first evaluated on the ImageNet validation dataset, with the same categories on which these two DCNNs were trained. Similar findings were revealed in the two DCNNs. First, the values of the PSI were low for all object categories in all layers in general (median < 0.4), with the maximum values no larger than 0.6 (Fig. 1), suggesting that the sparse coding scheme was broadly adopted in all layers of the DCNNs to represent objects. Interestingly, the degree of sparseness is comparable to 0.36 of neuronal population in the extrastriate cortex in macaques (Lehky et al., 2011). Second, in each layer, the PSI of all categories exhibited a broad distribution (ranges > 0.2), indicating great individual differences in sparseness among object categories. However, despite large amount of inter-category differences, the median PSI of each layer showed a trend of increase along the hierarchy in both Conv and FC layers respectively (AlexNet: Kendall’s tau = .40, *p* < .001; VGG11: Kendall’s tau = .36, *p* < .001). Note that the increase in sparseness was not strictly monotonic, as the PSI of the first layer was slightly higher than the adjacent ones. More interestingly, although AlexNet and VGG11 have different numbers of Conv layers, the major increase occurred at the last Conv layer.

**Figure 1.**
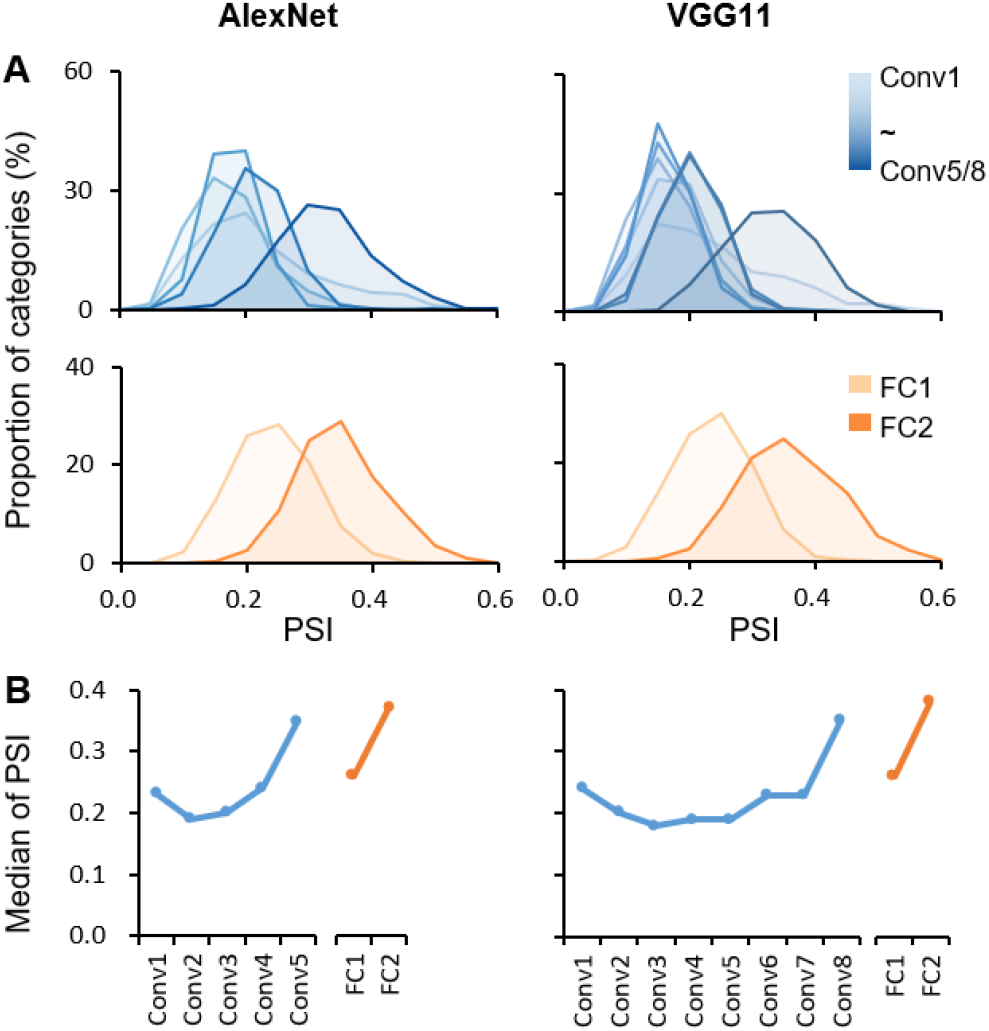
Hierarchically sparse coding for object categories in DCNNs. (**A)** Layer-wise PSI distribution for object categories in DCNNs. The sparseness was evaluated using the PSI for each object category from the ImageNet dataset (1,000 categories) in each layer separately. The distribution of PSI right-shifted along hierarchy in general. X axis: the degree of sparseness, with higher PSI indicating a higher degree of sparseness; Y axis: the proportion of categories with a corresponding PSI value. (**B)** Median of PSI for each layer. In general, the median of PSI increased along hierarchy in Conv and FC layers respectively. X axis: the name of layers along hierarchy; Y axis: the median of PSI.

We replicated this finding with a new dataset, Caltech256, that is dissimilar to the ImageNet in object categories and is thus not in the training dataset. We found a similar pattern of the increase in sparseness along the hierarchy (AlexNet: Kendall’s tau = .35, *p* < .001; VGG11: Kendall’s tau = .25, *p* < .001, supplemental Figure 1), suggesting that the increase in sparseness did not result from image dataset. Taken together, the hierarchically-increased sparseness suggested that there was a systematic shift from the distributed-like coding scheme in low layers to the local-like coding scheme in high layers.

Next, we examined the functionality of the sparse coding scheme observed in the DCNNs. To address this question, we tested the association between the population sparseness and the behavioral performance by performing correlation analyses within each layer of the DCNNs. In AlexNet, significant correlations were found starting from Conv4 and beyond [*r*s(254) > 0.19, *p*s < .001, Bonferroni corrected] (Fig. 2A). This result suggested that the degree of sparseness in coding object categories was predictive of performance accuracy. That is, the sparser an object category was represented, the better it was recognized and classified. Importantly, the correlation coefficients also increased along hierarchy (Kendall’s tau = .90, *p* = .003), with the highest correlation coefficient observed at Conv5 (0.43) and FC2 (0.69) respectively (Fig. 2A). This trend suggests a closer relationship between the population sparseness and the behavioral performance in higher layers. Indeed, with a stepwise multiple regression analysis in which PSI of all Conv/FC layers of certain categories were the independent variables and classification performance was the dependent variable, we confirmed that population sparseness was predictive of behavioral performance [Conv layers: F (3, 252) = 22.54, p < .001, adjusted R^2^ = 0.2; FC layers: F (2, 253) = 136.60, *p* < .001, adjusted R^2^ = 0.52; Fig. 2B]. Meanwhile, only PSI in higher layers starting from Conv3 remained in the regression models, further confirming that the coding scheme as a characteristic of representation became more essential with the increasing hierarchical level. Similar results were also found in VGG11 (Fig. 2B), suggesting that the association between sparseness and performance may be universal in DCNNs.

**Figure 2.**
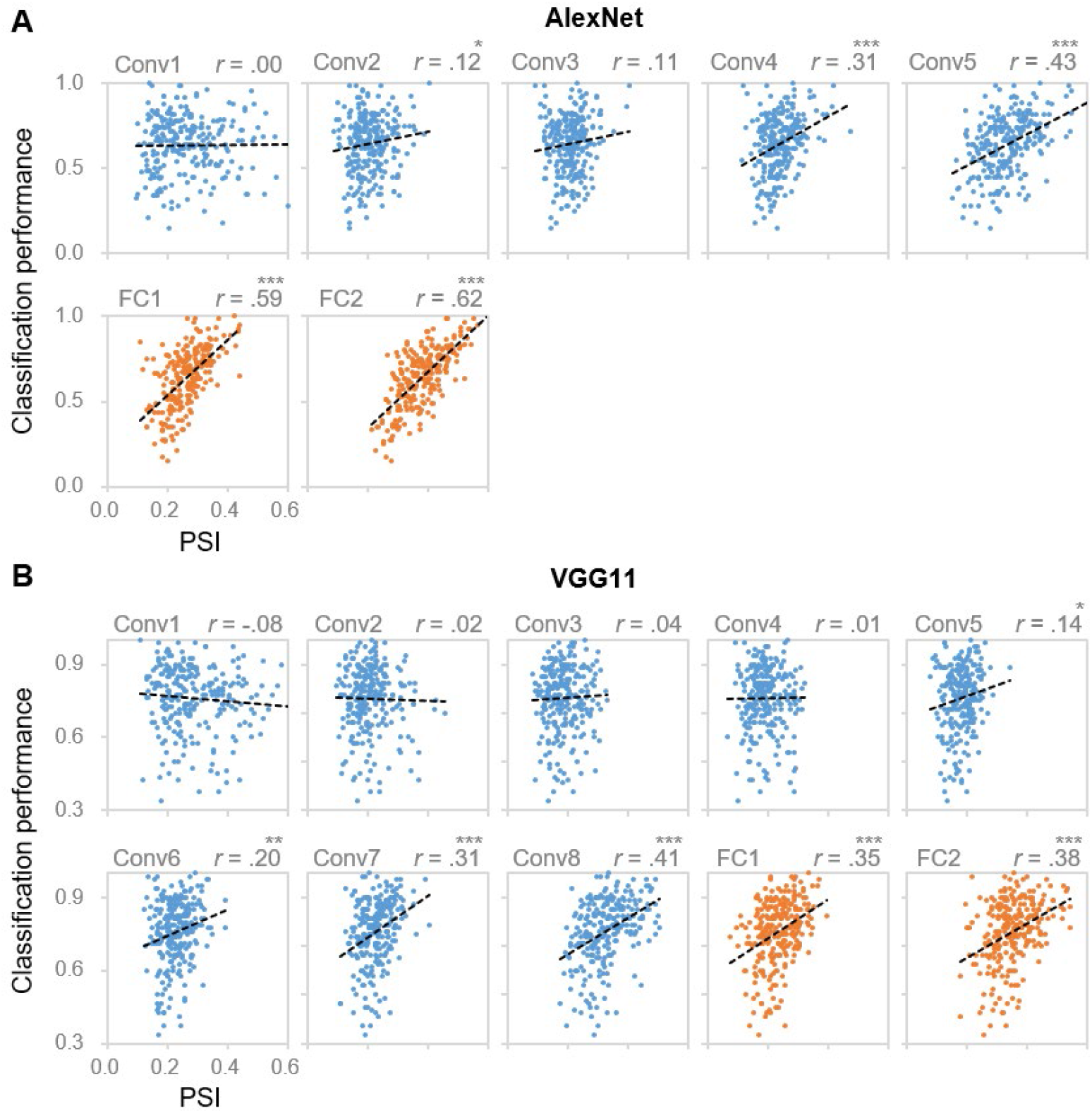
Correlation between coding sparseness and behavioral performance. Layer-wise scatter plots of DCNNs’ classification performance versus PSI values from **(A)** AlexNet and **(B)** VGG11 for object categories from Caltech256. X axis: PSI value, the larger the value the sparser the coding; Y axis: DCNNs’ classification performance for each object category. Each dot represents one category. * denotes *p* < .05, ** denotes *p* < .01 and *** denotes *p* < .001.

Finally, we explored the factors that may affect the formation of such a hierarchical coding scheme in the DCNNs. The DCNNs consist of two sublayers at a core in each layer (Fig. 3A), one is the feature extraction sublayer whose weights are dynamically adjusted during learning, and the other is a fixed nonlinear function (i.e., ReLU) that silences units with negative activities. To examine whether the hierarchically-increased sparseness was constructed through learning, we randomly permuted the weights of the pretrained AlexNet within each layer, and found that the degree of sparseness instead decreased along hierarchy (AlexNet: Kendall’s tau = −.57, *p* < .001; VGG11: Kendall’s tau = −.55, *p* < .001, Fig. 3B), which was contradictory to the finding of the undisrupted one (Fig. 1). In addition, when the ReLU sublayers were deactivated with the feature extraction sublayers intact (Fig. 3C), we also observed a decrease tendency of sparseness along the hierarchy (AlexNet: Kendall’s tau = −.21, *p* < .001; VGG11: Kendall’s tau = −.32, *p* < .001, Fig. 3D), again in contrast to the AlexNet with functioning ReLU (Fig. 1). Similar results were also found in VGG11, suggesting a general effect of learning and gating on the formation of the hierarchically-evolved coding scheme in DCNN.

**Figure 3.**
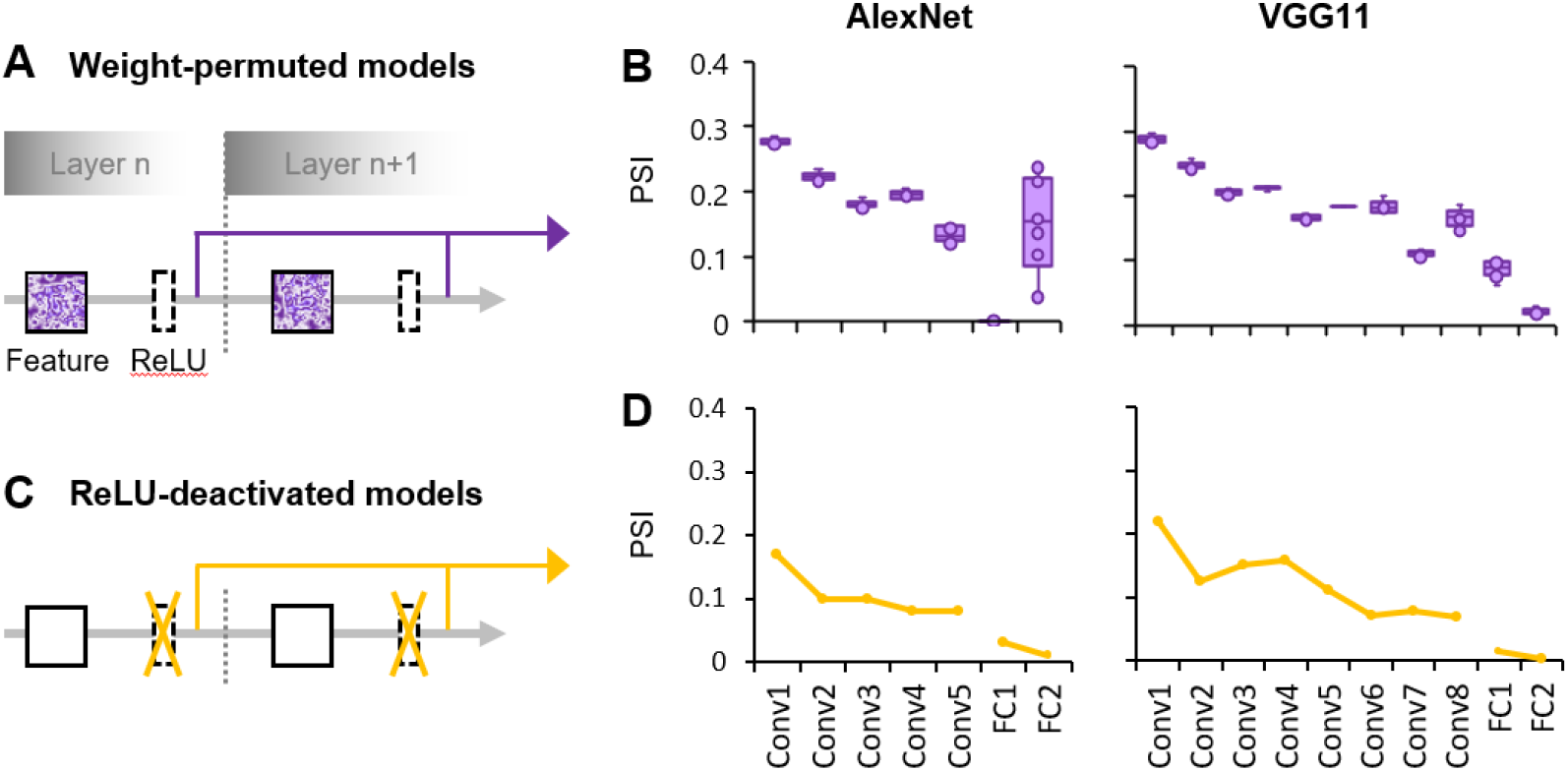
Both the learning process and the gating process play an important role in the formation of the hierarchically-evolved coding scheme in the DCNNs. **(A)** A schematic diagram of the weight-permuted models. **(B)** Box plots of median PSI for objects across layers in the weight-permuted models, which represent the minimum, maximum, median, first quartile and third quartile of the distribution of the median PSI values. The PSI was measured in 10 weight-permuted models using the same procedure as the intact one. **(C)** A schematic diagram of the ReLU-deactivated models. **(D)** Median PSI for objects across layers in the ReLU-deactivated models. X axis: the name of layers along hierarchy; Y axis: the median of PSI.

## 4 Discussion

In the present study, we systematically characterized the coding scheme in representing object categories at each layer of two typical DCNNs, AlexNet and VGG11. We found that objects were in general sparsely encoded in the DCNNs, and the degree of sparseness increased along the hierarchy. Importantly, the hierarchically-evolved sparseness was able to predict the classification performance of the DCNNs, revealing the functionality of the sparse coding. Finally, lesion analyses of the weight-permuted models and the ReLU-deactivated models suggest that the learning experience and the built-in gating operation account for the hierarchically sparse coding scheme in the DCNNs. In short, our study provided one of the first empirical evidence illustrating how object categories were represented in DCNNs for object recognition.

The finding that the degree of sparseness increased along the hierarchy in DCNNs is consistent with previous studies on DCNNs (Szegedy et al., 2013; Agrawal et al., 2014; Tripp, 2016; Wang et al., 2016; Morcos et al., 2018; Casper et al., 2019; Parde et al., 2020) and coincides with the long-time conjecture that the selectivity of neuronal responses to objects increases along the hierarchy in human and monkey’s visual ventral pathway (Perez-Orive, 2002; Norman and O’Reilly, 2003; Hung et al., 2005; Quiroga et al., 2005; Mormann et al., 2008). Our study further extended these previous studies by conducting a layer-wise analysis throughout all hierarchical levels and calculating the degree of sparseness based on responses of the entire population of units (‘neurons’ in DCNN). Besides, our study tested two datasets of more than 1,000 object categories, and thus provided more comprehensive coverage of the object space. Finally, we also examined the functionality of sparse coding by showing that the sparser an object category was encoded, the higher accuracy of the object category was correctly recognized. This type of study is impractical for studies on biological neural networks, because the number of objects, neurons and neural sites are largely limited by neurophysiological approaches, availability of subjects and ethical issues (Baddeley et al., 1997; Vinje and Gallant, 2000; Tolhurst et al., 2009). Taken together, our study not only systematically revealed the evolution of coding schemes along the hierarchy of artificial intelligent systems, but also suggests a perfect model to study mechanisms of biological intelligent systems.

The fact that both biological and artificial intelligent systems adopt a hierarchically-evolved sparse coding scheme suggests an implementation-independent principle of representing a myriad of objects efficiently. That is, at the lower level of vision, representations recruit a large number of generic units (i.e., distributed coding) to process natural objects with high fidelity. At the higher level, objects are decomposed and transformed into abstract features in the object space; therefore, only a smaller but highly-specialized units are needed to construct the representation. That is, a high degree of sparseness provides the DCNNs a more efficient way of processing objects without consuming too much energy (Lennie, 2003). Critically, a higher degree of sparseness also makes representations more interpretable, because only at higher layers the degree of sparseness was able to read out for behavioral performance. That is, the evolution of sparseness along the hierarchy likely mirrored the stages of objects being processed through the transformation of representation from stimulus-fidelity to goal-fidelity.

Interestingly, the sparseness was not accumulated gradually layer by layer. Instead, the sparseness was the highest at the last convolutional layer (i.e., Conv5 in AlexNet and Conv8 in VGG11) and fully-connected layer (i.e., FC2 in AlexNet and VGG11), much higher than that of their preceding ones regardless of the total number of layers in the DCNNs. This observation suggests a mechanism that the degree of sparseness dramatically increases at transitional layers either to the next processing stage (from Conv layers to FC layers) or to the generation of behavioral performance (from FC layers to the output layer). Further studies are needed to explore the functionality of the dramatical increase in sparseness.

As an intelligent system, DCNNs are a product of the predesigned architecture by nature and learned features by nurture. Our lesion study revealed that both architecture and learning were critical for the formation of the hierarchically sparse coding scheme. As for the innate architecture, a critical built-in function is the non-linear gating sublayer, ReLU, that silences neurons with negative activity (Glorot et al., 2011; LeCun et al., 2015). Obviously, the gating function is bound to increase the sparseness of coding because it removes weak or irrelevant activations and thus leads objects to be represented by a smaller number of units. Our study confirmed this intuition by showing the disruption of hierarchically-increased sparseness when the gating function being disabled. On the other hand, the gating function was not sufficient for a proper sparse coding scheme, because after randomly permuting the weights of the learned filters in the feature sublayers, the sparseness was no longer properly constructed either. Further, the dependence of both external learning experiences and built-in nonlinear operations implies that the sparse coding scheme may be also adopted in biological brains, because the gating function is the fundamental function of neurons (Lucas, 1909; Adrian, 1914) and the deprivation of visual experiences leads to deficits in a variety of visual functions (Wiesel and Hubel, 1963; Fine et al., 2003; Duffy and Livingstone, 2005).

In sum, our study on the coding scheme of object categories in DCNNs invites future studies to understand how in DCNN objects are recognized accurately in particular, and how intelligence emerges under the interaction of internal architecture and external learning experiences in general. On one hand, approaches and findings from neurophysiological and fMRI studies help to transpire the black-box of DCNNs and enlighten the design of more effective DCNNs. For example, our study suggests new algorithms for better performance by increasing sparseness effectively possibly through learning or gating function built in the network. On the other hand, in contrast to the fact that neurophysiological studies on nonhuman primates and fMRI studies on human are limited either by the coverage of brain areas or by the spatial resolution, both architecture and units’ activation in DCNNs are transparent. Therefore, DCNNs likely provides a perfect model to pry open mechanisms of object recognition at both micro- and macro-levels, which helps to understand how biological intelligent systems work.

## Acknowledgements

This study was funded by the National Natural Science Foundation of China (Grant No. 31861143039, 31771251), the National Key R&D Program of China (Grant No. 2019YFA0709503), and the National Basic Research Program of China (Grant No. 2018YFC0810602).

## Supplementary Material

**Supplementary Figure 1.**
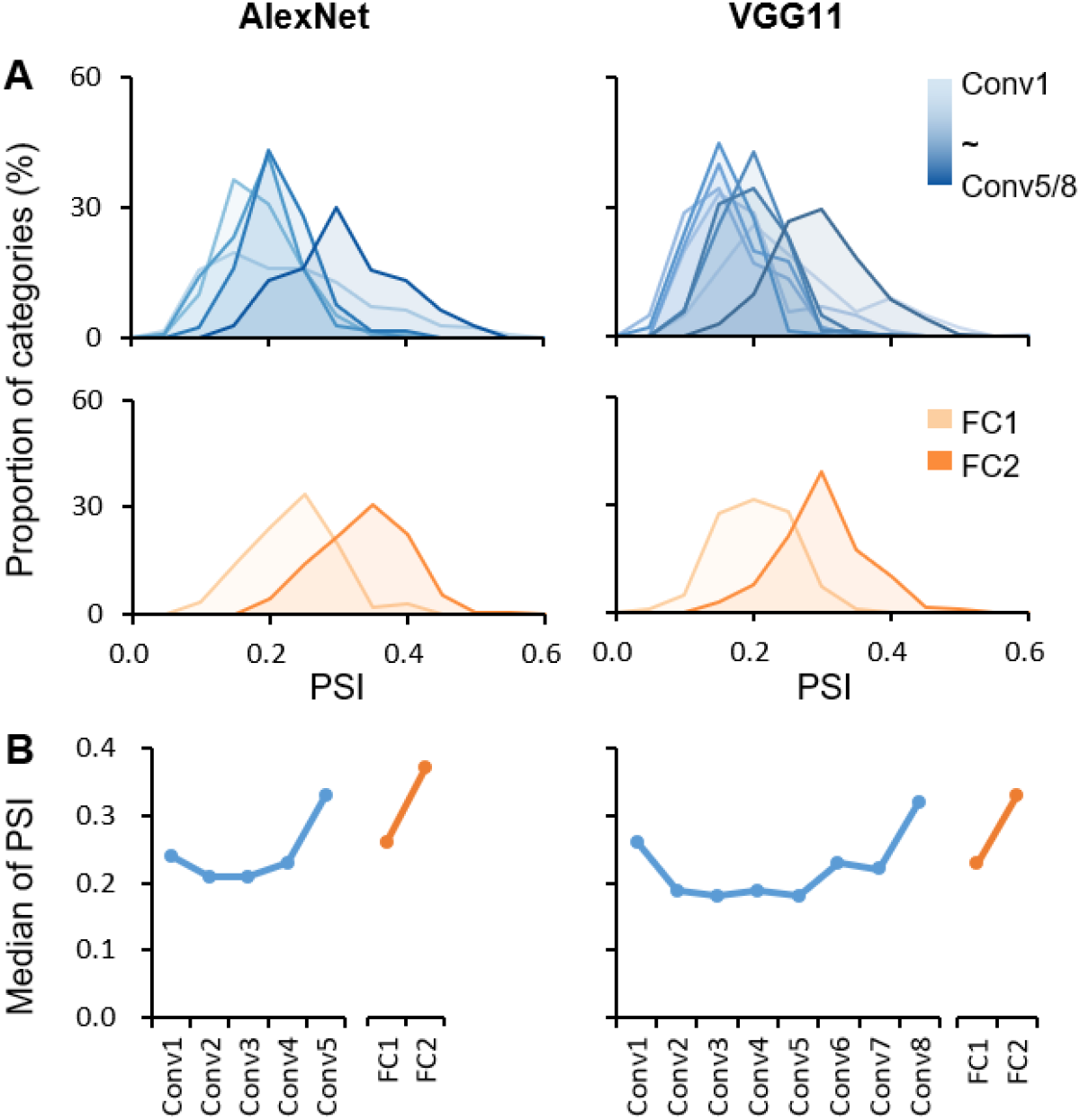
Hierarchically sparse coding for object categories in DCNNs on Caltech256 dataset. **(A)** Layer-wise PSI distribution for object categories in DCNNs on Caltech 256 dataset. Note that 113 categories among the 256 categories were excluded because of their overlap with the ImageNet dataset; therefore, the remaining 143 categories were used for validation. **(B)** Median of PSI for each layer. The median of PSI in general increased along Conv and FC layers respectively. A significant tendency was found for PSI across all layers (AlexNet: Kendall’s tau = .35, p < .001; VGG11: Kendall’s tau = .25, p < .001), which is consistent with the results found on ImageNet dataset as in Fig. 1.

## Notes

**Conflict of Interest:** None declared.

### Competing Interest Statement

The authors have declared no competing interest.

### Summary of Updates

Title in the file metadata revised

