## Supplemental Figure 1 for "Hierarchical sparse coding of objects in deep convolutional neural networks"

### Supplementary Material

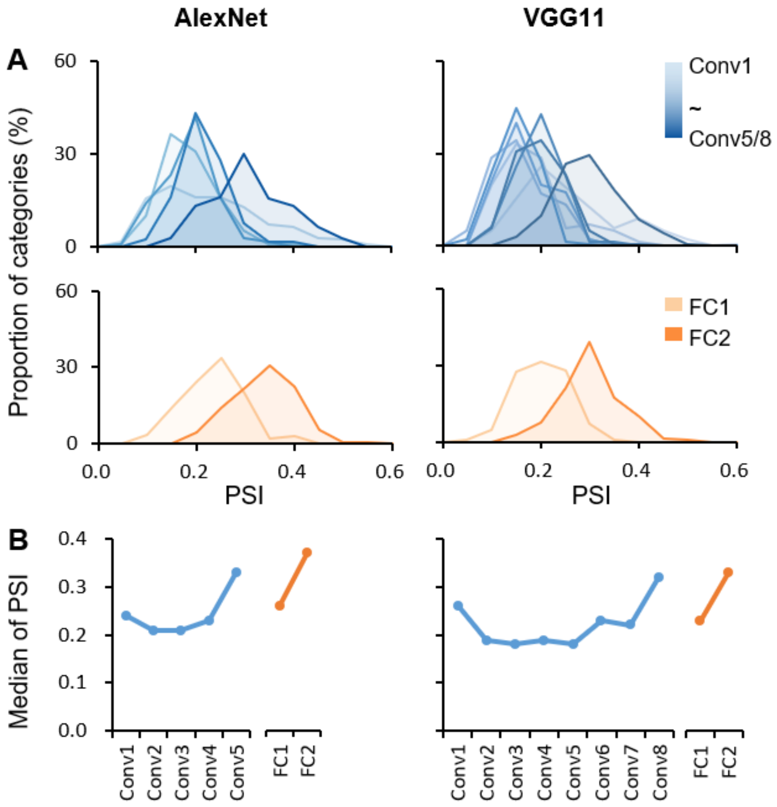


**Supplementary Figure 1. Hierarchically sparse coding for object categories in DCNNs on Caltech256 dataset. (A)** Layer-wise PSI distribution for objects in DCNNs on Caltech 256 dataset. Note that 113 categories among the 256 categories were excluded because of their overlap with the ImageNet dataset; therefore, the remaining 143 categories were used for validation. **(B)** Median of PSI for each layer. The median of PSI in general increased along Conv and FC layers respectively. A significant tendency was found for PSI across all layers (AlexNet: Kendall's tau = .35, $p < .001$; VGG11: Kendall's tau = .25, $p < .001$), which is consistent with the results found on ImageNet dataset as in Fig. 1.
